# Metacoder: An R Package for Visualization and Manipulation of Community Taxonomic Diversity Data

**DOI:** 10.1101/071019

**Authors:** Zachary S. L. Foster, Thomas J. Sharpton, Niklaus J. Grünwald

## Abstract

Community-level data, the type generated by an increasing number of metabarcoding studies, is often graphed as stacked bar charts or pie graphs that use color to represent taxa. These graph types do not convey the hierarchical structure of taxonomic classifications and are limited by the use of color for categories. As an alternative, we developed *metacoder*, an R package for easily parsing, manipulating, and graphing publication-ready plots of hierarchical data. *Metacoder* includes a dynamic and flexible function that can parse most text-based formats that contain taxonomic classifications, taxon names, taxon identifiers, or sequence identifiers. *Metacoder* can then subset, sample, and order this parsed data using a set of intuitive functions that take into account the hierarchical nature of the data. Finally, an extremely flexible plotting function enables quantitative representation of up to 4 arbitrary statistics simultaneously in a tree format by mapping statistics to the color and size of tree nodes and edges. *Metacoder* also allows exploration of barcode primer bias by integrating functions to run digital PCR. Although it has been designed for data from metabarcoding research, *metacoder* can easily be applied to any data that has a hierarchical component such as gene ontology or geographic location data. Our package complements currently available tools for community analysis and is provided open source with an extensive online user manual.

Note: This article was previously submitted as a pre-print: Zachary S. L. Foster, Thomas J. Sharpton, Niklaus J. Grünwald. 2016. *Metacoder*: An R package for manipulation and heat tree visualization of community taxonomic data from metabarcoding. BioRxiv 071019; doi: http://dx.doi.org/10.1101/071019.

## 1 Introduction

Metabarcoding is revolutionizing our understanding of complex ecosystems by circumventing the traditional limits of microbial diversity assessment, which include the need and bias of culturability, the effects of cryptic diversity, and the reliance on expert identification. Metabarcoding is a technique for determining community composition that typically involves extracting environmental DNA, amplifying a gene shared by a taxonomic group of interest using PCR, sequencing the amplicons, and comparing the sequences to reference databases [1]. It has been used extensively to explore communities inhabiting diverse environments, including oceans [2], plants [3], animals [4], humans [5], and soil [6].

The complex community data produced by metabarcoding is challenging conventional graphing techniques. Most often, bar charts, stacked bar charts, or pie graphs are employed that use color to represent a small number of taxa at the same rank (e.g. phylum, class, etc). This reliance on color for categorical information limits the number of taxa that can be effectively displayed, so most published figures only show results at a coarse taxonomic rank (e.g. class) or for only the most abundant taxa. These graphing techniques do not convey the hierarchical nature of taxonomic classifications, potentially obscuring patterns in unexplored taxonomic ranks that might be more biologically important. More recently, tree-based visualizations are becoming available as exemplified by the python-based MetaPhlAn and the corresponding graphing software GraPhlAn [7]. This tool allows visualization of high-quality circular representations of taxonomic trees.

Here, we introduce the R package *metacoder* that is specifically designed to address some of these problems in metabarcoding-based community ecology, focusing on parsing and manipulation of hierarchical data and community visualization in R. *Metacoder* provides a visualization that we call “heat trees” which quantitatively depicts statistics associated with taxa, such as abundance, using the color and size of nodes and edges in a taxonomic tree. These heat trees are useful for evaluating taxonomic coverage, barcode bias, or displaying differences in taxon abundance between communities. To import and manipulate data, *metacoder* provides a means of extracting and parsing taxonomic information from text-based formats (e.g. reference database FASTA headers) and an intuitive set of functions for subsetting, sampling, and rearranging taxonomic data. *Metacoder* also allows exploration of barcode primer bias by integrating digital PCR. All this functionality is made intuitive and user-friendly while still allowing extensive customization and flexibility. *Metacoder* can be applied to any data that can be organized hierarchically such as gene ontology or geographic location. *Metacoder* is an open source project available on CRAN and is provided with comprehensive online documentation including examples.

## 2 Design and Implementation

The R package *metacoder* provides a set of novel tools designed to parse, manipulate, and visualize community diversity data in a tree format using any taxonomic classification (Figure 1). Figure 1 illustrates the ease of use and flexibility of *metacoder*. It shows an example analysis extracting taxonomy from the 16S Ribosomal Database Project (RDP) training set for *mothur* [8], filtering and sampling the data by both taxon and sequence characteristics, running digital PCR, and graphing the proportion of sequences amplified for each taxon. Table 1 provides an overview of the core functions available in *metacoder*.

**Fig. 1.**
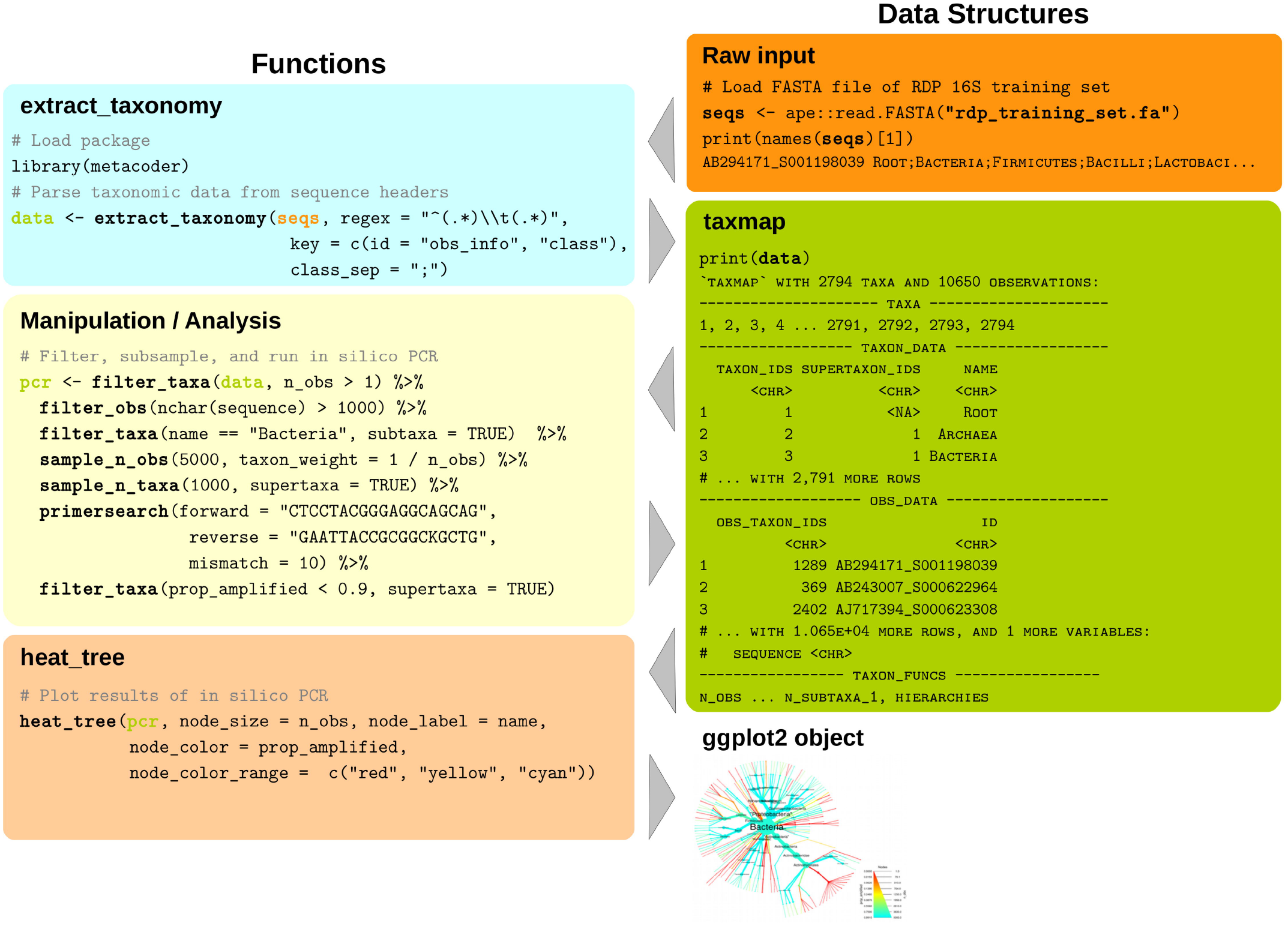
*Metacoder* has an intuitive and easy to use syntax. The code in this example analysis parses the taxonomic data associated with sequences from the Ribosomal Database Project [9] 16S training set, filters and subsamples the data by sequence and taxon characteristics, conducts digital PCR, and displays the results as a heat tree. All functions in bold are from the *metacoder* package. Note how columns and functions in the `taxmap` object (green box) can be referenced within functions as if they were independent variables.

### 2.1 The taxmap data object

To store the taxonomic hierarchy and associated observations (e.g. sequences) we developed a new data object class called `taxmap.` The `taxmap` class is designed to be as flexible and easily manipulated as possible. The only assumption made about the users data is that it can be represented as a set of observations assigned to a hierarchy; the hierarchy and the observations do not need to be biological. The class contains two tables in which user data is stored: a taxonomic hierarchy stored as an edge list of unique IDs and a set of observations mapped to that hierarchy (Figure 1). Users can add, remove, or reorder both columns and rows in either `taxmap` table using convenient functions included in the package (Table 1). For each table, there is also a list of functions stored with the class that each create a temporary column with the same name when referenced by one of the manipulation or plotting functions. These are useful for attributes that must be updated when the data is subset or otherwise modified, such as the number of observations for each taxon (see “n_obs” in Figure 1). If this kind of derived information was stored in a static column, the user would have to update the column each time the data set is subset, potentially leading to mistakes if this is not done. There are many of these column-generating functions included by default, but the user can easily add their own by adding a function that takes a `taxmap` object. The names of columns or column-generating functions in either table of a `taxmap` object can be referenced as if they were independent variables in most *metacoder* functions in the style of popular R packages like *ggplot2* and *dplyr*. This makes the code much easier to read and write.

### 2.2 Universal parsing and retrieval of taxonomic information

*Metacoder* provides a way to extract taxonomic information from text-based formats so it can be manipulated within R. One of the most inefficient steps in bioinformatics can be loading and parsing data into a standardized form that is usable for computational analysis. Many databases have unique taxonomy formats with differing types of taxonomic information. The structure and nomenclature of the taxonomy used can be unique to the database or reference another database such as GenBank [10]. Rather than creating a parser for each data format, *metacoder* provides a single function to parse any format definable by regular expressions that contains taxonomic information (Figure 1). This makes it easier to use multiple data sources with the same downstream analysis.

The `extract_taxonomy` function can parse hierarchical classifications or retrieve classifications from online databases using taxon names, taxon IDs, or Genbank sequence IDs. The user supplies a regular expression with capture groups (parentheses) and a corresponding key to define what parts of the input can provide classification information. The `extract_taxonomy` function has been used successfully to parse several major database formats including Genbank [10], UNITE [11], Protist Ribosomal Reference Database (PR2) [12], Greengenes [13], Silva [14], and, as illustrated in figure 1, the RDP [9]. Examples for each database are provided in the user manuals [15].

**Table 1:**
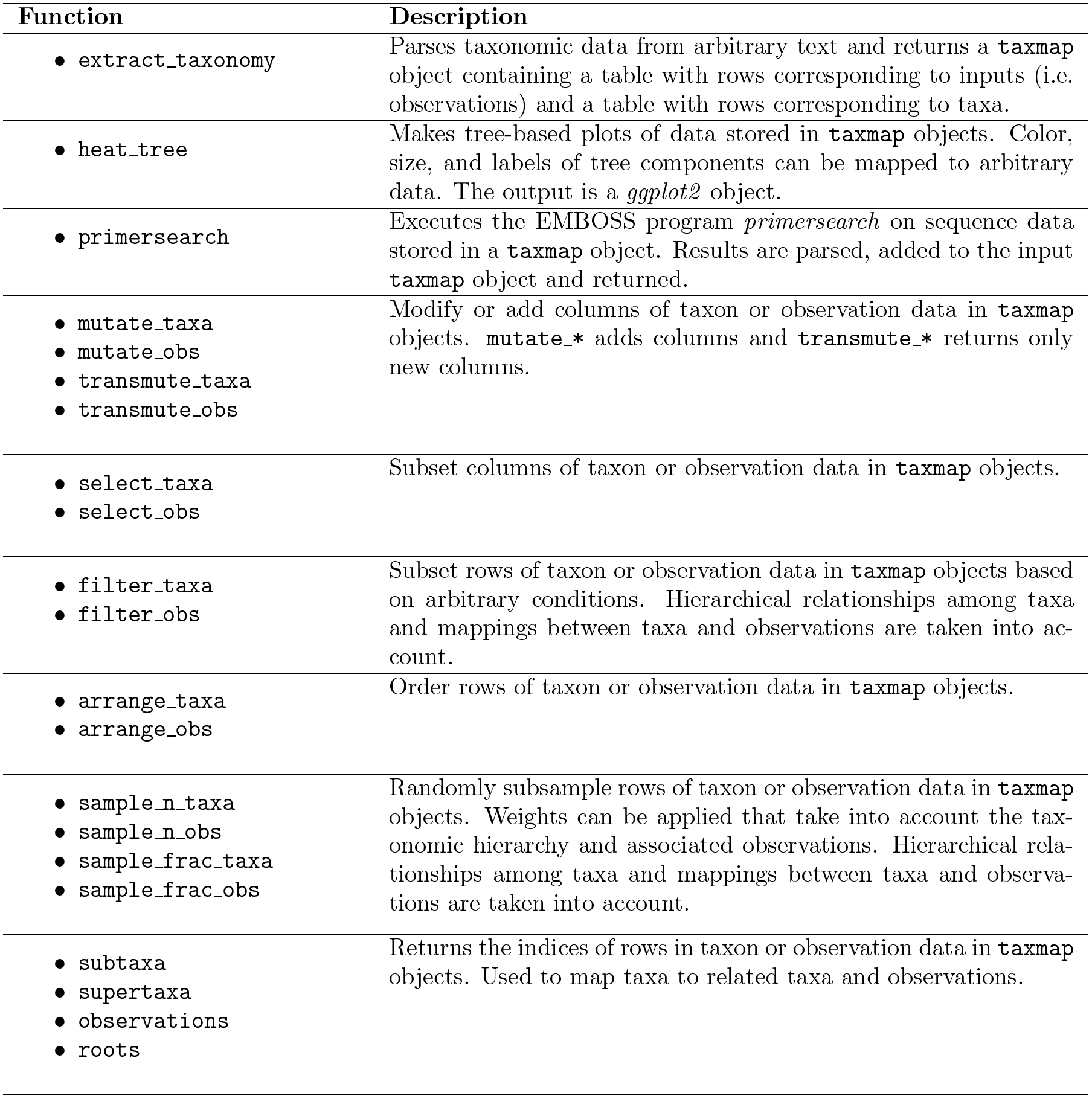
Primary functions found in *metacoder.*

### 2.3 Intuitive manipulation of taxonomic data

*Metacoder* makes it easy to subset and sample large data sets composed of thousands of observations (e.g. sequences) assigned to thousands of taxa, while taking into account hierarchical relationships. This allows for exploration and analysis of manageable subsets of a large data set. Taxonomies are inherently hierarchical, making them difficult to subset and sample intuitively compared with typical tabular data. In addition to the taxonomy itself, there is usually also data assigned to taxa in the taxonomy, which we refer to as “observations”. Subsetting either the taxonomy or the associated observations, depending on the goal, might require subsetting both to keep them in sync. For example, if a set of taxa are removed or left out of a random subsample, should the subtaxa and associated observations also be removed, left as is, or reassigned to a supertaxon? If observations are removed, should the taxa they were assigned to also be removed? The functions provided by *metacoder* gives the user control over these details and simplifies their implementation.

*Metacoder* allows users to intuitively and efficiently subset complex hierarchical data sets using a cohesive set of functions inspired by the popular *dplyr* data-manipulation philosophy. *Dplyr* is an R package for providing a conceptually consistent set of operations for manipulating tabular information [16]. Whereas *dplyr* functions each act on a single table, *metacoder*’s analogous functions act on both the taxon and observation tables in a `taxmap` object (Table 1). For each major *dplyr* function there are two analogous *metacoder* functions: one that manipulates the taxon table and one that manipulates the observations table. The functions take into account the relationship between the two tables and can modify both depending on parameterization, allowing for operations on taxa to affect their corresponding observations and vice versa. They also take into account the hierarchical nature of the taxon table. For example, the *metacoder* functions `filter_taxa` and `filter_obs` are based on the *dplyr* function `filter` and are used to remove rows in the taxon and observation tables corresponding to some criterion. Unlike simply applying a filter to these tables directly, these functions allow the subtaxa, supertaxa, and/or observations of taxa passing the filter to be preserved or discarded, making it easy to subset the data in diverse ways (Figure 1). There are also functions for ordering rows (`arrange_taxa, arrange_obs),` subsetting columns `(select_taxa, select_obs),` and adding columns `(mutate_taxa, mutate_obs).`

*Metacoder* also provides functions for random sampling of taxa and corresponding observations. The function `taxonomic_sample` is used to randomly sub-sample items such that all taxa of one or more given ranks have some specified number of observations representing them. Taxa with too few sequences are excluded and taxa with too many are randomly subsampled. Whole taxa can also be sampled based on the number of sub-taxa they have. Alternatively, there are *dplyr* analogues called `sample_n_taxa` and `sample_n_obs,` which can sample some number of taxa or observations. In both functions, weights can be assigned to taxa or observations, influencing how likely each is to be sampled. For example, the probability of sampling a given observation can be determined by a taxon characteristic, such as the number of observations assigned to that taxon, or it could be determined by an observation characteristic, like sequence length. Similar to the `filter_*` functions, there are parameters controlling whether selected taxa’s subtaxa, supertaxa, or observations are included or not in the sample (Figure 1).

### 2.4 Heat tree plotting of taxonomic data

Visualizing the massive data sets being generated by modern sequencing of complex ecosystems is typically done using traditional stacked barcharts or pie graphs, but these ignore the hierarchical nature of taxonomic classifications and their reliance on colors for categories limits the number of taxa that can be distinguished (Figure 2). Generic trees can convey a taxonomic hierarchy, but displaying how statistics are distributed throughout the tree, including internal taxa, is difficult. *Metacoder* provides a function that plots up to 4 statistics on a tree with quantitative legends by automatically mapping any set of numbers to the color and width of nodes and edges. The size and content of edge and node labels can also be mapped to custom values. These publication-quality graphs provide a method for visualizing community data that is richer than is currently possible with stacked bar charts. Although there are other R packages that can plot variables on trees, like *phyloseq* [17], these have been designed for phylogenetic rather than taxonomic trees and therefore optimized for plotting information on the tips of the tree and not on internal nodes.

**Fig. 2.**
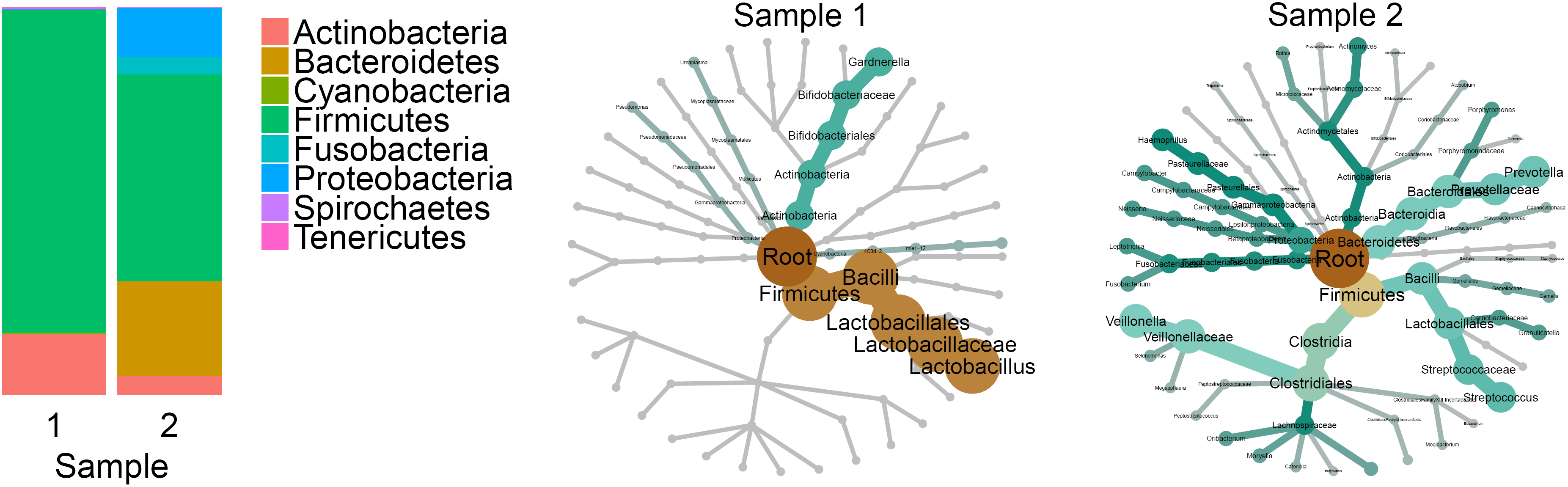
Heat trees allow for a better understanding of community structure than stacked bar charts. The stacked bar chart on the left represents the abundance of organisms in two samples from the Human Microbiome Project [5]. The same data are displayed as heat trees on the right. In the heat trees, size and color of nodes and edges are correlated with the abundance of organisms in each community. Both visualizations show communities dominated by firmicutes, but the heat trees reveal that the two samples share no families within firmicutes and are thus much more different than suggested by the stacked bar chart.

The function `heat_tree` creates a tree utilizing color and size to display taxon statistics (e.g., sequence abundance) for many taxa and ranks in one intuitive graph (Figure 2). Taxa are represented as nodes and both color and size are used to represent any statistic associated with taxa, such as abundance. Although the `heat_tree` function has many options to customize the appearance of the graph, it is designed to minimize the amount of user-defined parameters necessary to create an effective visualization. The size range of graph elements is optimized for each graph to minimize overlap and maximize size range. Raw statistics are automatically translated to size and color and a legend is added to display the relationship. Unlike most other plotting functions in R, the plot looks the same regardless of output size, allowing the graph to be saved at any size or used in complex, composite figures without changing parameters. These characteristics allow `heat_tree` to be used effectively in pipelines and with minimal parameterization since a small set of parameters displays diverse taxonomy data. The output of the `heat_tree` function is a *ggplot2* object, making it compatible with many existing R tools. Another novel feature of heat trees is the automatic plotting of multiple trees when there are multiple “roots” to the hierarchy. This can happen when, for example, there are “Bacteria” and “Eukaryota” taxa without a unifying “Life” taxon, or when coarse taxonomic ranks are removed to aid in the visualization of large data sets (Figure 3).

**Fig. 3.**
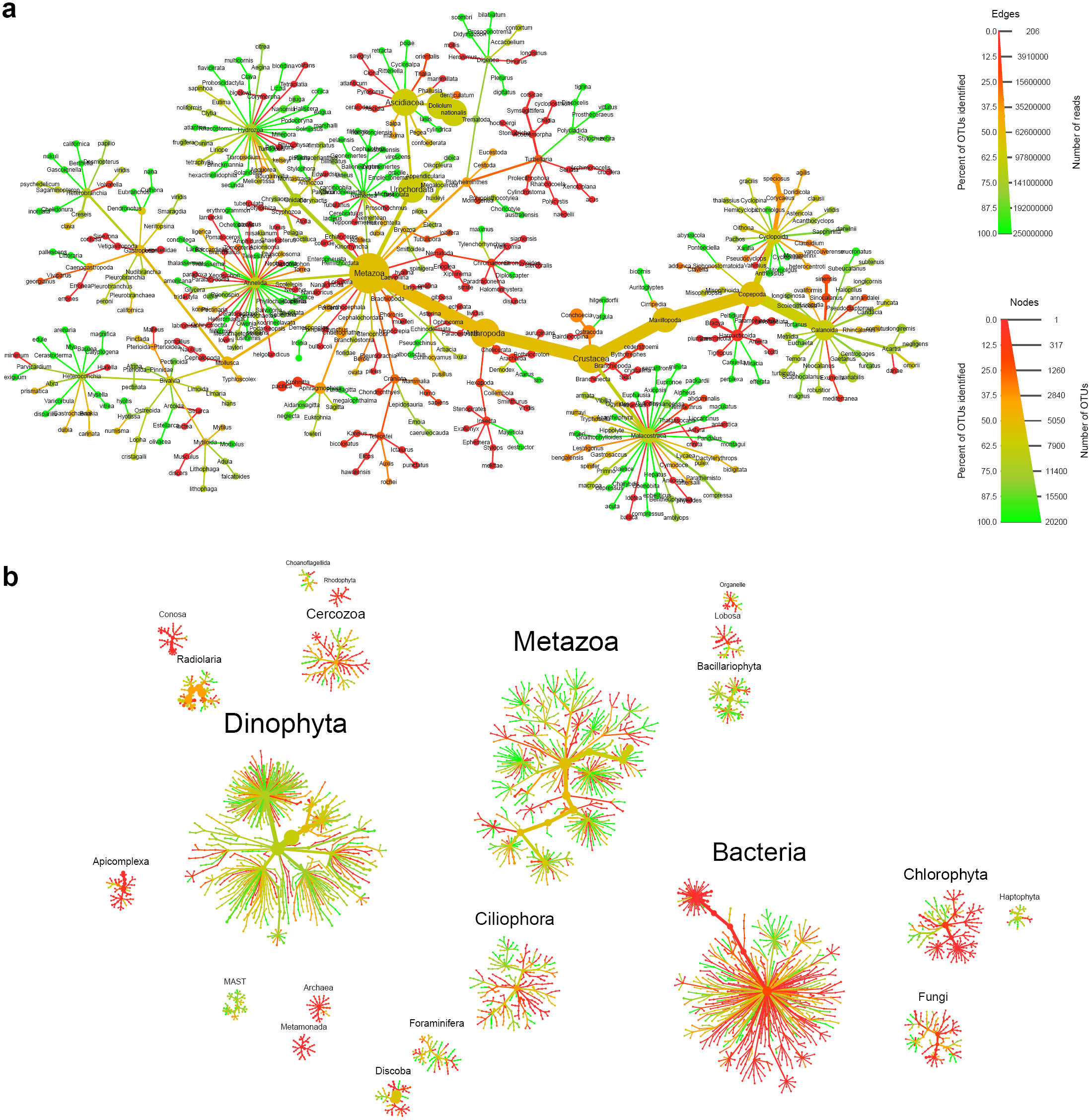
Heat trees display up to four metrics in a taxonomic context and can plot multiple trees per graph. Most graph components, such as the size and color of text, nodes, and edges, can be automatically mapped to arbitrary numbers, allowing for a quantitative representation of multiple statistics simultaneously. This graph depicts the uncertainty of OTU classifications from the TARA global oceans survey [2]. Each node represents a taxon used to classify OTUs and the edges determine where it fits in the overall taxonomic hierarchy. Node diameter is proportional to the number of OTUs classified as that taxon and edge width is proportional to the number of reads. Color represents the percent of OTUs assigned to each taxon that are somewhat similar to their closest reference sequence (>90% sequence identity). `a.` Metazoan diversity in detail. `b.` All taxonomic diversity found. Note that multiple trees are automatically created and arranged when there are multiple roots to the taxonomy.

## 3 Results

### 3.1 Heat trees allow quantitative visualization of community diversity data

We developed heat trees to allow visualization of community data in a taxonomic context by mapping any statistic to the color or size of tree components. Here, we reanalyzed data set 5 from the TARA oceans eukaryotic plankton diversity study to visualize the similarity between OTUs observed in the data set and their closest match to a sequence in a reference database [2]. The TARA ocean expedition analyzed DNA extracted from ocean water throughout the world. Even though a custom reference database was made using curated 18S sequences spanning all known eukaryotic diversity, many of the OTUs observed had no close match. Figure 3 shows a heat tree that illustrates the proportion of OTUs that were well characterized in each taxon (at least 90% identical to a reference sequence). Color indicates the percentage of OTUs that are well characterized, node width indicates the number of OTUs assigned to each taxon, and edge width indicates the number of reads. Taxa with ambiguous names and those with less than 200 reads have been filtered out for clarity. This figure illustrates one of the principal advantages of heat trees, as it reveals many clades in the tree that contain only red lineages, which indicate that the entire taxonomic group is poorly represented in the reference sequence database. Of particular interest are those clades with predominantly red lineages that also have relatively large nodes, such as Harpacticoida (in Copepoda on the right). These represent taxonomic groups that were found to have high amounts of diversity in the oceans, but for which we have a paucity of genomic information. Investigators interested in improving the genomic resolution of the biosphere can thus use these approaches to rapidly assess which taxa should be prioritized for focused investigations. Note that a large portion of the taxa shown in red, yellow or orange have many OTUs with a poor match to the reference taxonomic hierarchy.

### 3.2 Flexible parsing allows for similar use of diverse data

Metabarcoding studies often rely on techniques or data that may introduce bias into an investigation. For example, the specific set of PCR primers used to amplify genomic DNA and the taxonomic annotation database can both have an effect on the study results. A quick and inexpensive way to estimate biases caused by primers is to use digital PCR, which simulates PCR success using alignments between reference sequences and primers. *Metacoder* can be used to explore different databases or primer combinations to assess these effects since it supplies functions to parse divserse data sources, conduct digital PCR, and plot the results. Figure 4 shows a series of heat tree comparisons that were produced using a common 16S rRNA metabarcoding primer set [18] and digital PCR against the full-length 16S sequences found in three taxonomic annotation databases: Greengenes [13], RDP [9], and SILVA [14]. These heat trees reveal subsets of the full taxonomies for these three databases that poorly amplify by digital PCR using the selected primers. As a result, they indicate which lineages within each of the taxonomies may be challenging to detect in a metabarcoding study that uses these primers. Importantly, different sets of primers likely amplify different sets of taxa, so investigators interested in specific lineages can use this approach in conjunction with various primer sets to identify those that maximize the likelihood of discovery and reduce wasted sequencing resources on non-target organisms. However, these heat maps do not indicate whether one database is necessarily preferable over another, as they differ in the structure of their taxonomies, as well as the number and phylogenetic diversity of their reference sequences. For example, most of the bacterial clades that do not amplify well in the SILVA lineages are unnamed lineages that are not found in the other databases, indicating that they warrant further exploration.

**Fig. 4.**
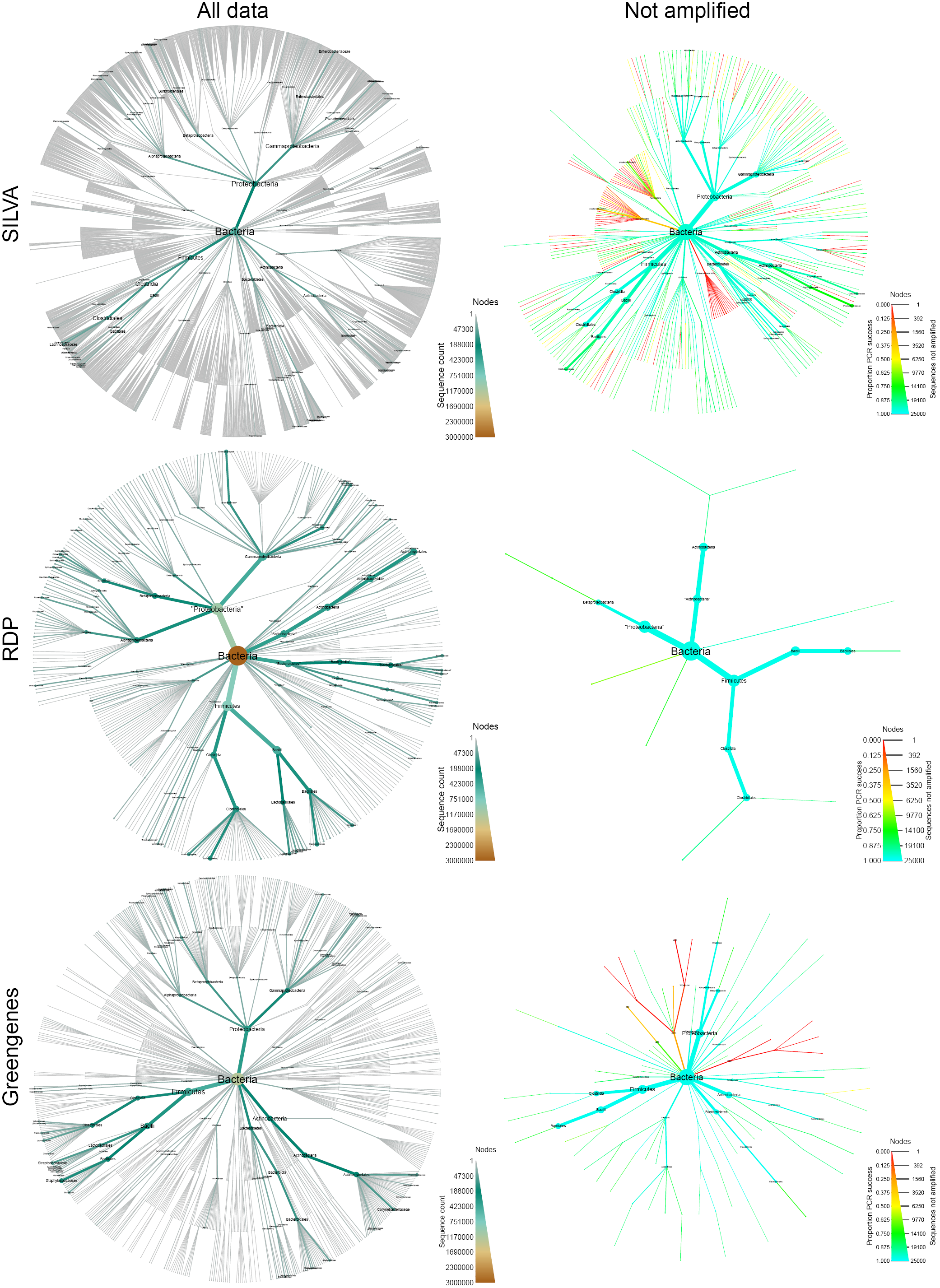
Flexible parsing and digital PCR allows for comparisons of primers and databases. Shown is a comparison of digital PCR results for three 16S reference databases. The plots on the left display abundance of all bacterial 16S sequences. Plots on the right display all taxa with subtaxa not entirely amplified by digital PCR using universal 16S primers [19]. Node color and size display the proportion and number of sequences not amplified respectively.

### 3.3 Heat trees can show pairwise comparisons of communities across treatments

One challenge in metabarcoding studies is visually determining how specific sub-sets of samples vary in their taxonomic composition. Unlike most other graphing software in R, *metacoder* produces graphs that look the same at any output size or aspect ratio, allowing heat trees to be easily integrated into larger composite figures without changing the code for individual subplots. Using color to depict the difference in read or OTU abundance between two treatments can result in particularly effective visualizations, especially when the presence of color is made dependent on a statistical test. To examine more than two treatments at once, a matrix of these kind of heat trees can be combined with a labeled “guide” tree. Figure 5 shows application of this idea to human microbiome data showing pairwise differences between body sites. Coloring indicates significant differences between the median proportion of reads for samples from different body sites as determined using a Wilcox rank-sum test followed by a Benjamini-Hochberg (FDR) correction for multiple testing. The intensity of the color is relative to the log-2 ratio of difference in median proportions. Brown taxa indicate an enrichment in body sites listed on the top of the graph and green is the opposite. While the original study [5] showed abundance plots, our visualization provides the taxonomic context. For example, *Haemophilus, Streptococcus*, and *Prevotella* spp. are enriched in saliva (brown) relative to stool where *Bacteroides* is enriched (green). We also see that in the Lachnospiraceae clade several genera shown in both green and brown taxa are differentially abundant. These observations are consistent with known differences in the human-associated microbiome across body sites, but heat trees uniquely provide an integrated view of how all levels of a taxonomy vary for all pairs of body sites.

**Fig. 5.**
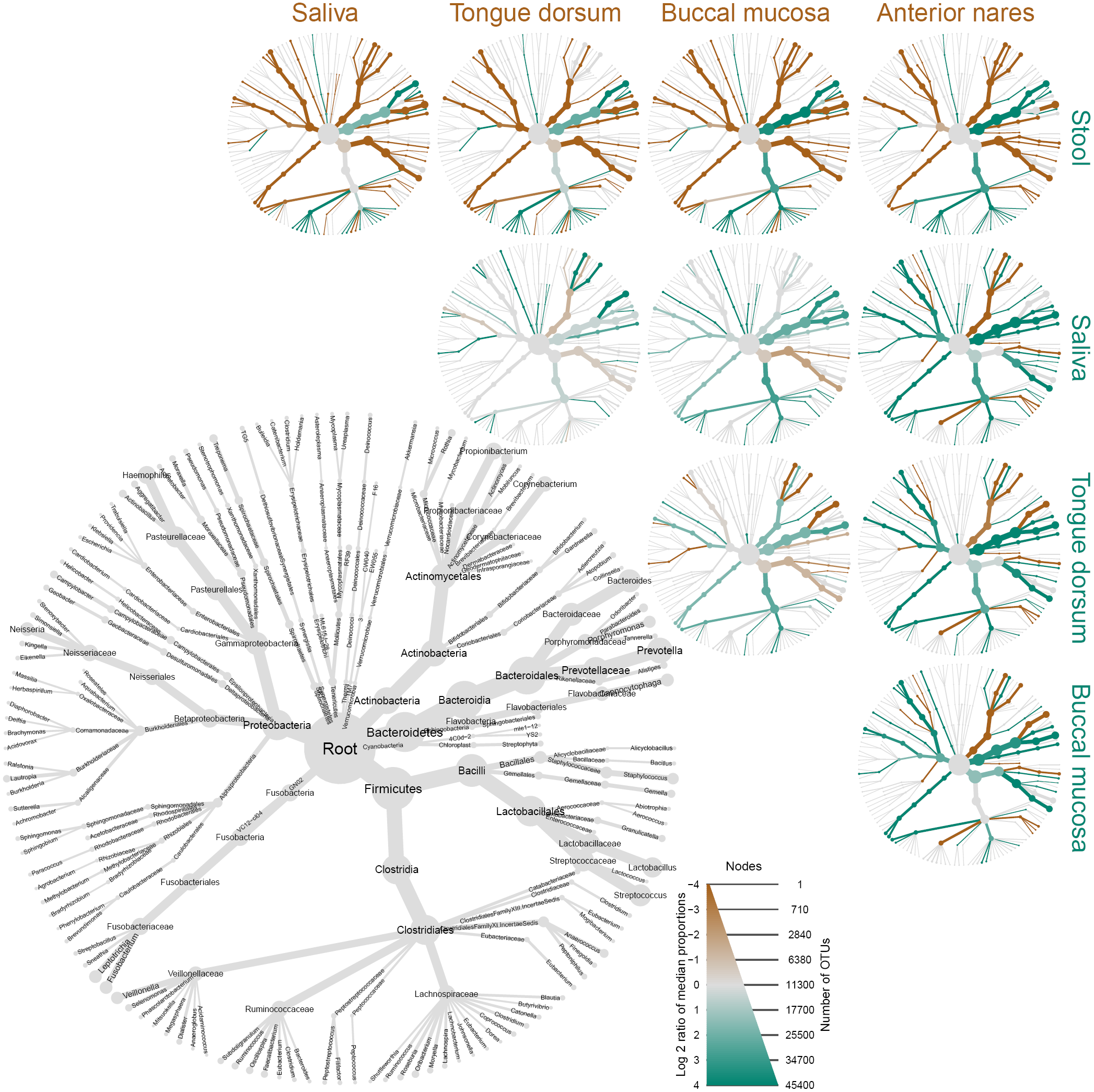
Scale-independent appearance facilitates complex, composite figures. All graph components, including text, have the same relative sizes independent of output size, unlike most graphical packages in R, making it easier to create composite figures entirely within R. This graph uses 16S metabarcoding data from the human microbiome project study. The gray tree on the lower left functions as a key for the smaller unlabeled trees. The color of each taxon represents the log-2 ratio of median proportions of reads observed at each body site. Only significant differences are colored, determined using a Wilcox rank-sum test followed by a Benjamini-Hochberg (FDR) correction for multiple comparisons. For example, *Haemophilus, Streptococcus, Prevotella* are enriched in saliva (brown) relative to stool where *Bacteroides* is enriched (green).

### 3.4 Other applications

The `taxmap` data object defined in *metacoder* can be used for any data that can be classified by a hierarchy. Figure 6, for example, shows an analysis of votes cast in the 2016 US Democratic party national primaries organized by geography. The heat tree reveals distinct patterns such as a sweep by Clinton in the South and a split on the West coast, with California predominantly voting for Clinton while Washington and Oregon predominantly voted for Sanders. Another potential application is displaying the results of gene expression studies by associating differential expression with gene ontology (GO) annotations. Figure 7 shows the results of a RNA-seq study on the effect of glucocorticoids on smooth muscle tissue [20]. All biological processes influenced by at least one gene with a significant change in expression are plotted. The authors of the study find that genes involved in immune response are influenced by the glucocorticoid treatment. Viewing these results in a heat tree shows not only the specific immune process affected (the branch on the middle right), but also the more general phenomena they constitute; regulation of high level phenomena, like immune system function, can be explained by specific processes like “leukocyte chemotaxis” and these specific processes are put into the context of the phenomena they contribute to. This is more informative than simply reporting the results for a single level of the GO annotation hierarchy or discussing the effects of genes one at a time.

**Fig. 6.**
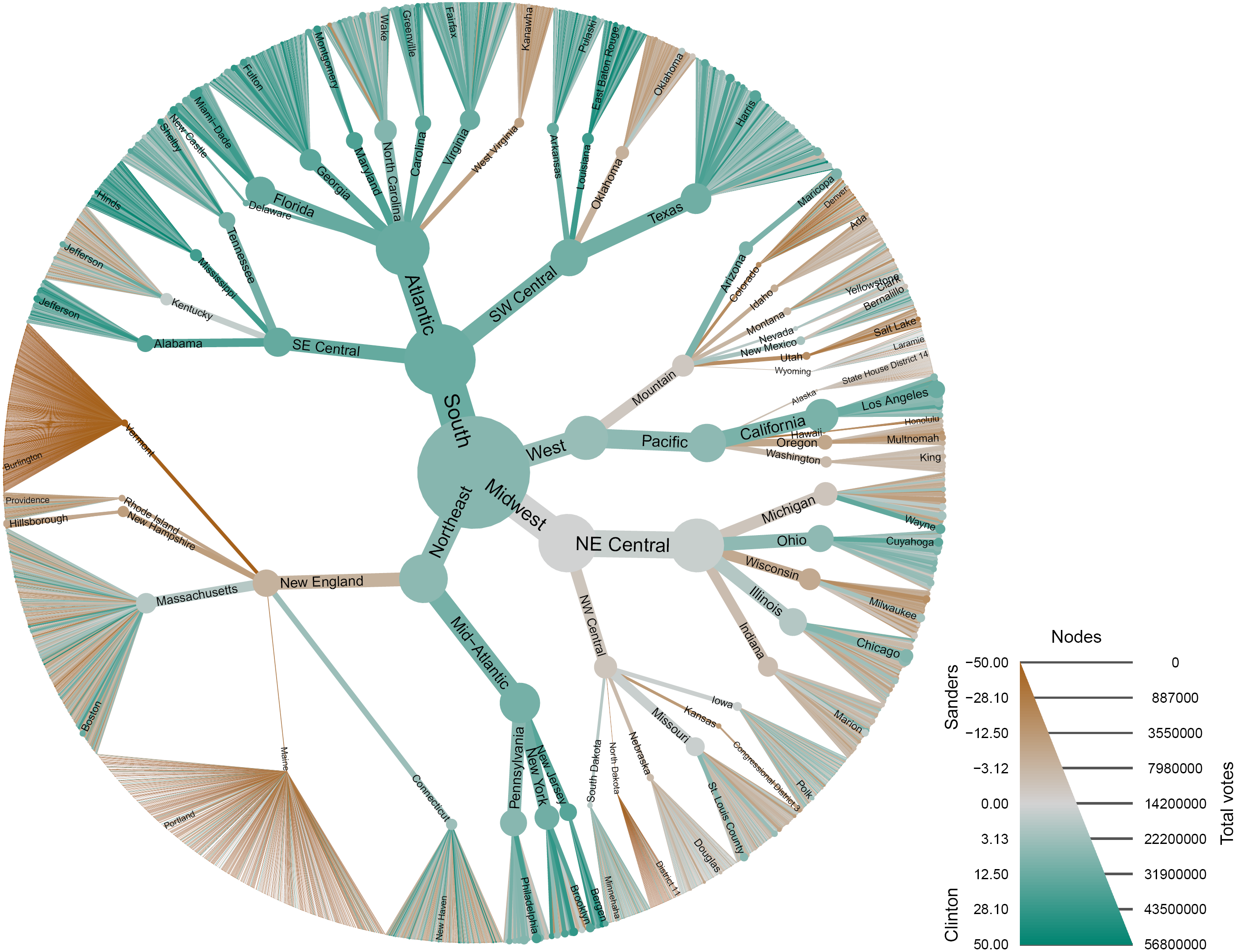
*Metacoder* can be used with any type of data that can be organized hierarchically. This plot shows the results of the 2016 Democratic primary election organized by region, division, state, and county. The regions and divisions are those defined by the United States census bureau. Color corresponds to the difference in the percentage of votes for candidates Hillary Clinton (green) and Bernie Sanders (brown). Size corresponds to the total number of votes cast. Data was downloaded from https://www.kaggle.com/benhamner/2016-us-election/.

**Fig. 7.**
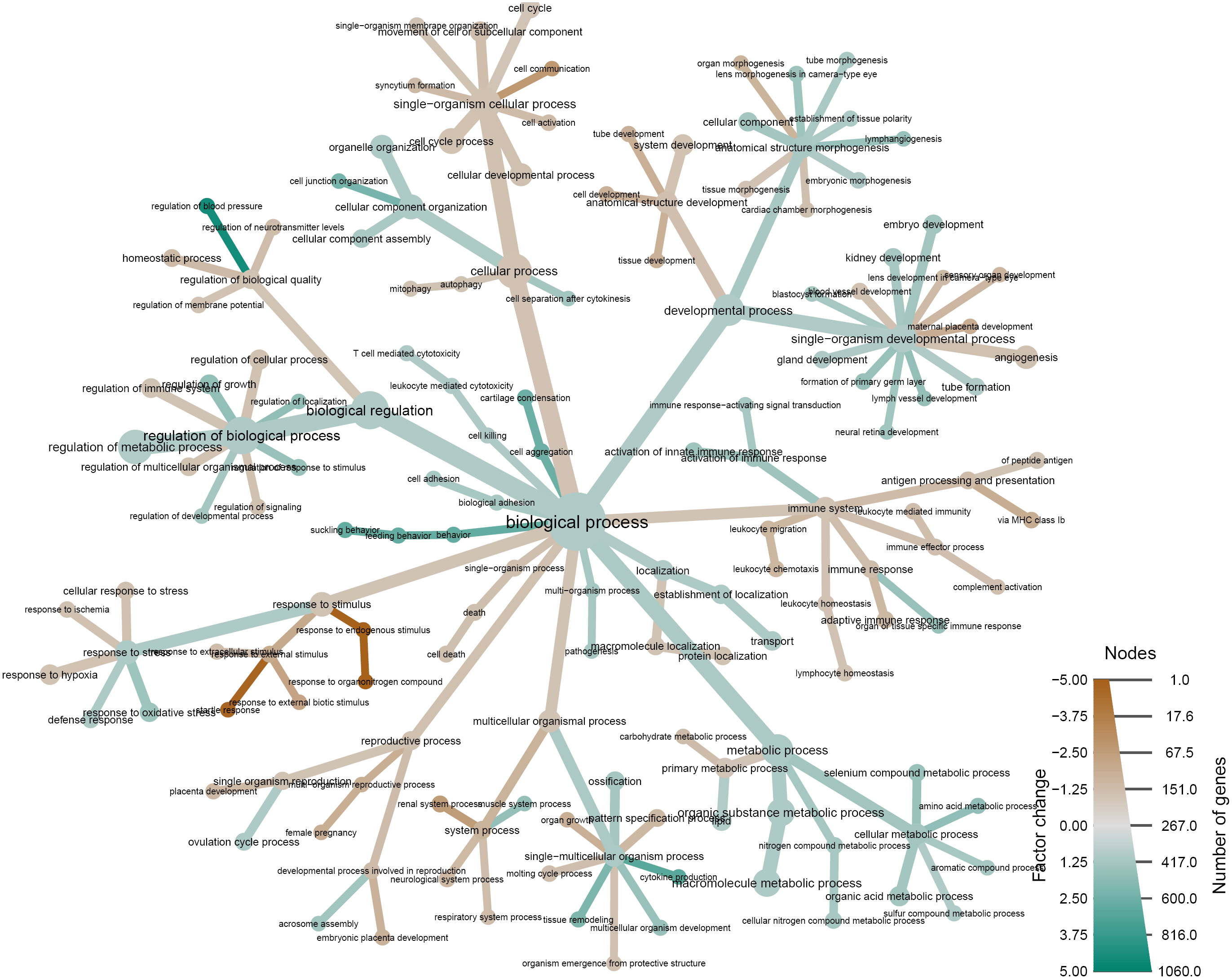
Another alternate use example: vizualizing gene expression data in a GO hierarchy. The gene ontology for all differentially expressed genes in a study on the effect of a glucocorticoid on airway smooth muscle tissue [20]. Color indicates the sign and intensity of averaged changes in gene expression and the size indicates the number of genes classified by a given gene ontology term.

## 4 Availability and Future Directions

The R package *metacoder* is an open-source project under the MIT License. Stable releases of *metacoder* are available on CRAN while recent improvements can be downloaded from github (https://github.com/grunwaldlab/metacoder). A manual with documentation and examples is provided [15]. This manual also provides the code to reproduce all figures included in this manuscript.

We are currently continuing development of *metacoder*. We welcome contributions and feedback from the community. We want to make *metacoder* functions and classes compatible with those from other bioinformatic R packages such as *phyloseq, ape, seqinr*, and *taxize*. We might integrate more options for digital PCR and barcode gap analysis, perhaps using ecoPCR or the R packages *PrimerMiner* and *Spider*. We are also considering adding additional visualization functions.

## 5 Acknowledgments

This work was supported in part by funds from USDA ARS CRIS Project 2027-22000-039-00 and the USDA ARS Floriculture Nursery Research Initiative. The use of trade, firm, or corporation names in this publication is for the information and convenience of the reader. Such use does not constitute an official endorsement or approval by the United States Department of Agriculture or the Agricultural Research Service of any product or service to the exclusion of others that may be suitable.

## 6 Author Contributions

Conceived and designed the experiments: ZSLF, NJG, TJS. Performed the experiments: ZSLF. Analyzed the data: ZSLF. Contributed reagents/materials/analysis tools: ZSLF, NJG. Wrote the paper: ZSLF, NJG, TJS. Designed, developed scripts: ZSLF.

